# Predicting Residues Involved in Anti-DNA Autoantibodies with Limited Neural Networks

**DOI:** 10.1101/2020.08.06.240101

**Authors:** Rachel St.Clair, Michael Teti, Mirjana Pavlovic, William Hahn, Elan Barenholtz

**Affiliations:** Florida Atlantic University, Center for Complex Systems and Brain Sciences, Boca Raton, Florida 33431 United States; Florida Atlantic University, Department of Electrical Engineering and Computer Science, Boca Raton, Florida 33431 United States

**Keywords:** Auto-Immunity, Deep Learning, DNA-Binding, LSTM, Proteomics, Systemic Lupus

## Abstract

Computer-aided rational vaccine design (RVD) and synthetic pharmacology are rapidly developing fields that leverage existing datasets for developing compounds of interest. Computational proteomics utilizes algorithms and models to probe proteins for functional prediction. A potentially strong target for such a computational approach is autoimmune antibodies which are the result of broken tolerance in the immune system where it cannot distinguish “self” from “non-self” resulting in attack of its own structures (proteins and DNA, mainly). The information on structure, function and pathogenicity of autoantibodies may assist in engineering RVD against autoimmune diseases. Current computational approaches exploit large datasets curated with extensive domain knowledge, most of which include the need for many computational resources and have been applied indirectly to problems of interest for DNA, RNA, and monomer protein binding. Here, we present a novel method for discovering potential binding sites. We employed long short-term memory (LSTM) models trained on FASTA primary sequences directly to predict protein binding in DNA-binding hydrolytic antibodies (abzymes). We also employed CNN models applied to the same dataset. While the CNN model outperformed the LSTM on the primary task of binding prediction, analysis of internal model representations of both models showed that the LSTM models highlighted sub-sequences that were more strongly correlated with sites known to be involved in binding. These results demonstrate that analysis of internal processes of recurrent neural network models may serve as a powerful tool for primary sequence analysis.

## 1 Introduction

Computational proteomics utilizes algorithms and models to probe proteins for functional prediction. Primary research in this area is often devoted to computer-aided rational vaccine design (RVD) and synthetic pharmacology for effective drug design. A potentially strong target for such a computational approach is autoimmune antibodies, which are the result of broken tolerance in the immune system where it cannot distinguish “self” from “non-self” resulting in attack of its own structures (proteins and DNA, mainly). Despite decades of research, much remains poorly understood about the the mechanisms underlying autoantibody function and binding processes. Computational approaches may represent a novel avenue for discovery, leading to the development of RVD for autoimmune diseases.

Considered to be a hallmark of lupus disease, anti-DNA antibody is found in 70-90% of patients with SLE (particularly in those with nephritis), and measurements of its levels in patients’ plasma is used to follow the course of disease. However, because anti-DNA antibody has been shown to be both hydrolytic and nephritogenic in a limited number of experimental and clinical studies, and that it also appears before the flare, it is suggested that it may serve as a strong flare predictor [1–3]. The important role of anti-DNA antibody is supported by studies in mouse models of nephritogenic lupus in which anti-DNA antibodies were found [4] as well as by the findings of [5] and [6]. The chemical structure and processes underlying autoantibodies remain poorly understood. [4,7,8] isolated anti-DNA and confirmed their DNA catalytic activities. However, only a small number of anti-DNA bindng antibodies’ binding sites have been determined. Almost an entire decade of X-ray crystallographic studies performed by [9] combined with the most recent data generated by [10–12] observed that tyrosine and tryptophan residues create a hydrophobic pocket within the side chain of the antibody [13,14]. Thus, Oligo-Thymidine pentamer enters the hydrophobic pocket between Tyrosine and Tryptophane from anti-DNA autoantibody in Fab fragment; where they bind to DNA, starting hydrolytic cleavage as a newly known modality of activity in autoimmune pathology (abzyme activity).

Wet-lab sequencing and X-ray crystallography are costly and time consuming, requiring expertise on each particular antibody. Computational approaches, which model existing data to generate novel predictions, can serve to narrow the field of possible candidates that may then be lab tested. Such models have become a standard tool in -omics research, with significant contributions to synthetic protein design and discovery. Recently, deep learning models have far exceeded earlier computational methods in complex feature detection from large datasets, as in the Large Scale Visual Recognition Challenge (ILSVRC) and machine generated text models like GPT-2 [15,16]. The unique ability of deep learning networks to define and manipulate important nonlinear features allows the possibility for such models to provide more insightful context than wet-lab and other traditional methods could alone. In recent years, deep learning has been applied to many areas within computational proteomics including protein folding, subcellular localization, and binding motif prediction, classification, and detection [17–19]. Indeed, nearly all recent computational approaches involve state of the art machine-learning including natural language processing (NLP) techniques, such as encoder-decoder networks and Recurrent Neural Networks (RNNs), Support Vector Machines (SVM), Convolutional Neural Networks (CNNs), and use-case specific optimization algorithms, etc. [20–25].

Most approaches to computational proteomics to date are heavily dependent on hand annotated datasets, supplementary feature input, require extensive background information, and/or are most frequently applied to large generic datasets. It is often the case in novel fields of interest that only limited, smaller datasets, lacking extra domain knowledge (ie. evolutionary, MSA, tertiary structure data) beyond primary sequence, are available. To date, only a handful of studies have applied machine learning to primary sequence alone to perform protein class —though not binding site— prediction [26,27]. With respect to binding site prediction, DNA and RNA specificities have been achieved using CNN, RNN, and hand-tailored MSA algorithms to other datasets (namely TFB and RNAB proteins) by [20–22]. However, these studies used both microarray and sequencing data. Most recently, [28, 29] achieved moderate accuracy in protein-protein interaction interface residue pairs prediction, but used supplementary data and hand-tailored algorithms for inference.

Thus to date, no computational studies have reported successful bindingsite prediction from primary sequence alone. As noted, in the case of most novel domains of interest without supplementary domain knowledge, a model capable of analysing primary sequence alone would be highly useful. Here we introduce a novel approach to achieve this goal based on analysis of hidden activation weights in RNNs, a family of deep learning models that include a form of memory, making them well adapted to analyzing sequential data. In particular, we used a class of RNNs — Long Short-Term Memory networks, or LSTMs – which include a memory cell to represent long-term memory allowing for sequential feature detection of position-specific input arrays [30].

LSTMs are well adapted to learning from biological sequences, such as proteins, because of their ability to analyze sequences at multiple levels including a whole protein, residue-to-residue interactions, or individual amino acids. More specifically, LSTMs are suited for the binding domain problem because proteins, presented as primary sequence, can be evaluated beyond their linear representation for features across the primary space that might signify binding in later tertiary structures.

### 1.1 Current Approach

Here, we trained several LSTM-based models to classify antibody protein primary sequences as DNA-binding or non-binding and then evaluated the model’s hidden-states to assess the potential of specific sub-sequences and residues as binding sites. We designed our deep learning model to be fully compatible with the protein data warehouse Uniprot [31]. For comparison, we also trained several CNN-based models of similar complexity on the same data. [27] used a similar LSTM model to predict phylogenetically distinct protein families by sequence alone and again in [29] predicted residue specificities but required more than just primary sequence data. Our work takes a similar approach in model architecture, making use of as few parameters as possible from only primary sequences, for the anti-DNA antibody problem set which is lacking in the amount of biological data available. We directly apply this model to further the implications of the hydrolytic activity exerted on DNA by autoantibodies of various length and phylogenetically distinct protein families (e.g. IGG, IGM, etc.). We assessed the applicability of our technique to a small, unascertained problem set to directly elucidate the anti-DNA autoantibody phenomena in a way that allows insight into the model’s inference process. To our knowledge, this work is the first application of small LSTM and CNN models to predict residues related to binding function from primary sequence alone and is the first computational model of anti-DNA antibodies.

In both LSTM and CNN cases, we use two models of different sized trainable parameters to predict binding from primary sequences. We evaluated each of the models with regard to binding prediction accuracy. In addition, we evaluated the subsequences indicated by the hidden activations of the different model for agreement with previously literature identified binding sites.

## 2 Related Work

Although x-ray crystallography is capable of elucidating the DNA binding domain in an antibody, it is typically expensive and time consuming since only one highly reliable protein can be processed at a time. Research in this area, spanning the last several decades, has not achieved the goal of a comprehensive understanding of the antibody binding motif involved in DNA recognition and later hydrolytic activity. Most extensive wet-lab work has been completed by [10–12] for a few proteins, both synthetic and de novo. DNA-binding motif prediction has been achieved in several early works using MSA [32], physics-based simulations [33], and kernel based algorithms [34]. Deep learning based approaches most often include the using of RNNs as in [23,24,26–29,35]. Some other recent works include combinations of CNN and RNN models [20–22]. All of these studies depended on large datasets, supplementary data, and/or millions of model parameters. Some recent work suggests that that protein primary sequence may not be sufficiently high-dimensional enough for the successful application of deep-learning techniques [36], which may account for the lack of sequence-only approaches in the literature. Furthermore, in [20] DNA/RNA position specific sites for TFB proteins were extracted using a brute-force approach by mutating each possible codon in areas of interest, determined by deep learning models, and accessing the respective binding score. This type of approach suffers from combinatorial explosion when applied to protein binding-sites as there are 27 possible residues (instead of 4 codons). This problem is exacerbated by variable protein length, which can often reach 2000 residues in length and shown to be problematic in the most similar works by [29]. The advantage of antecedent works are their ability to analyze giant datasets and high fidelity in their own applications. However, these works cannot be easily adapted to the problem presented in this work and others like it due to the limited domain information available, haven’t been shown reliable for sequence-only analysis, and completely forgo hidden state interpretation.

## 3 Methods

### 3.1 Dataset

An anti-DNA antibody dataset was curated directly from the protein data warehouse, Uniprot.org, using the query keywords: ‘Immunoglobuline’ and ‘DNA-binding’ in the manually annotated and reviewed records. This method supplied primary sequences of around 780 DNA binding related antibodies. The counter class was sourced the same way with the exclusionary keyword ‘NOT+DNA-binding’, which resulted in 1,267 antibodies. The data was first inspected for basic discrepancies between binding and non-binding antibodies. Each dataset was brute-force searched for amino acid frequency and length.

The generated dataset was found to include proteins of MHC and T-cell type that are not antibodies. This reflects the fact that Uniprot pulls all proteins associated with a keyword, but are not exclusive; meaning, the sequences originally retrieved relate to antibody function but may not be antibodies themselves. To create an unambiguous class of antibodies, we queried the generated data removing any proteins associated with MHC and T-cell keywords. After excluding these, only 75 antibody DNA-binding proteins remained, Therefore, we collected extra samples manually from the Protein DataBank website [37] using keywords, ‘DNA-binding’ and ‘Antibodies’. 33 test sequences were hand-selected according to relevance and reliability. After removing duplicates and sequences with lengths less than 50 or greater than 2000 amino acids long, among all datasets, 81 binding antibodies were retrieved. To downsample the non-binding class into a generally representative dataset, we use principal component analysis (PCA) on multiple randomly selected samples of 81 proteins until the PCA more closely resembled the bind sequences’ PCA [Fig. 3.1]. Close resemblance was determined by the operator upon visual examination. This dataset was split into training and validation by randomly sampling and checking for sequence length balance. Finally, this secondary dataset consisted of 61 sequences reserved for training and 20 for validation for the LSTM and CNN binding inference.

**Fig. 1.**
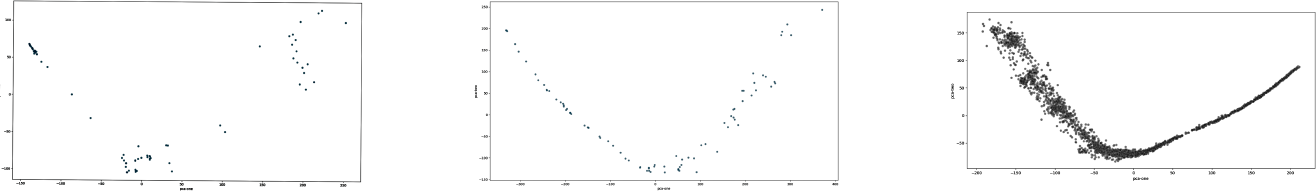
Raw Sequence PCA. Binding dataset PCA (left) and Non-bind dataset PCA after (middle) and before downsampling (right).

### 3.2 Pre-Processing

As a preliminary processing step, t-distributed stochastic neighbor embedding (TSNE), a technique for nonlinear clustering, was performed on the original 2,047 sequences to determine if features existed in the dataset that would lend to easy determination of class.

### 3.3 Data Augmentation

This secondary dataset was converted to one-hot images and augmented in two ways. As an LSTM evaluates a sequence, it uses recurrent information to update the hidden state and make classification decisions. The hyper-variable domain (HVD), Fab fragment, is most likely to be involved in ligand binding recognition in antibodies and is often written first in FASTA sequences. Since the hidden state is lacking recurrent information in the beginning of each sequence analysis, the hidden state values are often much higher than later time-steps in the data image. Therefore, to preserve the hidden state’s attention to the important HVD, we reversed all sequences. Data was then augmented with horizontal flips to increase the amount of data available since the one-hot encoding was arbitrarily created from left to right. Augmentation did not significantly increase model performance, but may have increased the robustness of the hidden state evaluation.

### 3.4 Binding Prediction

#### 3-4-1 LSTM Prediction

With a total of 244 training sequences and 80 validation sequences, the model was trained with one LSTM layer of 300 hidden nodes, a 50% dropout layer, and a 2-node fully connected layer for 200 epochs with a batch size of one [Fig. 2]. This model has 395,402 trainable parameters. Adam and cross-entropy loss were used as the criterion and optimizers for the model parameters. Due to variability in accuracy caused by random weight initialization and random batch sampling during training, 100 models were trained. The same process was repeated for a smaller LSTM with only 200 hidden nodes incurring 183,602 parameters. We later observed an increase in LSTM prediction accuracy with the addition of a final sigmoid activation and thus included it in both small and large LSTM model variants. Model weights were saved according to their best validation accuracy scores for later hidden state extraction.

**Fig. 2.**
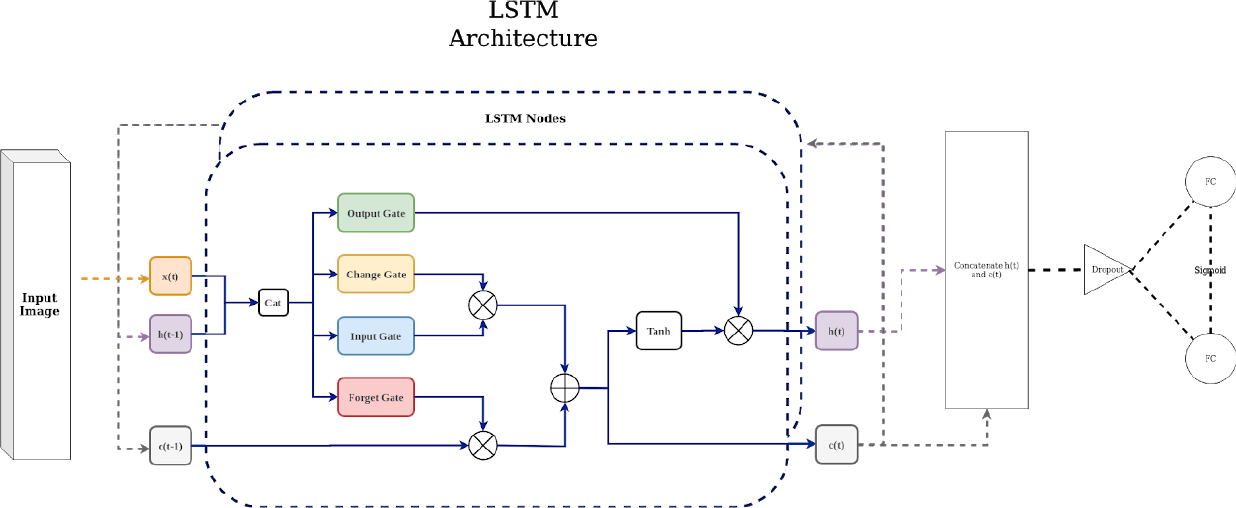
LSTM Model Architecture.

#### 3.4.2 CNN Prediction

The CNN was designed to have a similar number of parameters (394,425) as the LSTM network in order to equate the models to the greatest extent possible. Sequences were encoded as one-hot grayscale images, padded with an arbitrary value to the maximum sequence length of 1,750 and sampled with a batch size of one. The model consisted of three convolutional layers each followed by dropout and ReLU. The network’s final linear layer output 2 nodes and was evaluated by cross-entropy loss. The last convolutional layer was designed to retain the input sequences’ size outputting one feature map of size 1,751 by 28. A smaller variation of the model with only 183,743 parameters was also evaluated. [Fig. 3]. Hidden states were extracted from the best performing CNNs by summing the last convolutional layer’s feature map across all sequences and then across the one-hot encoding dimension. This allowed for total activation for each position to be calculated and processed similarly to the LSTM hidden state analysis. Best performing models were selected according to the same procedure as used in the LSTM binding-site analysis method.

**Fig. 3.**
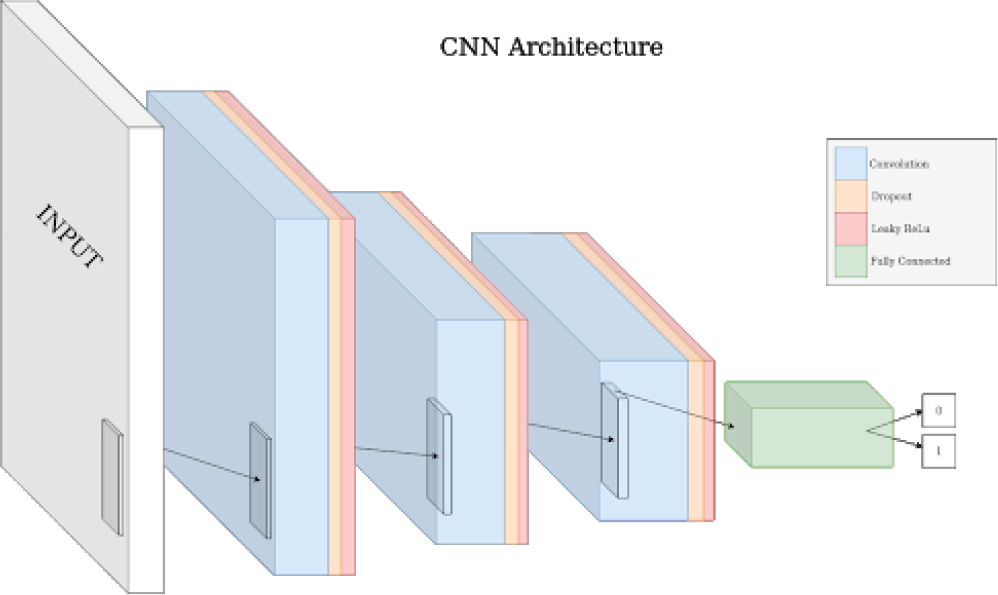
CNN Model Architecture.

### 3.5 Binding-site Analysis

Evaluation of the LSTM states’ hidden layer and CNN feature map activations were performed by extracting the respective weights from all correctly predicted, reversed sequences at each time-step for the top five performing models from each model variant. Since negative weights don’t necessarily mean negation in class prediction, the absolute value of all hidden cell activation weights (LSTM) and last convolutional layer (CNN) feature maps were recovered. Top models were those that most accurately predicted all sequences during testing of all 81 sequences in each class using the previously trained models’ learned weights. Once the best performing test models were determined, their original training and validation loss and accuracy trends were evaluated for obvious overfitting (i.e. poor training accuracy, loss in validation lower than loss in accuracy, etc.). All hidden cell weights were reversed so positions now align with FASTA formatting (position 0 is the first residue in the sequence and so forth). PCA was performed on the hidden cell weights of all top models combined for LSTM and CNN. Then, all sequence weights were summed per class, per model. Differences in position-specific areas of interest were first visualized in all weights for each time-step across the summed weight matrices. Activation weights were then summed across all nodes per class and scaled between 0 and 1 for comparison. The following tests were performed collectively on the five top models for each of the four model variants.

### 3.6 DNA-1 anti-DNA autoantibody

Activation weights for DNA binding antibody DNA-1 were recovered individually and compared to the position-specific residues important for binding given by x-ray crystallography reported in (Tanner, 2001). To remove the models’ internal representations of non-binding proteins and make activations more interpretable, convolutions were performed with 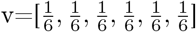 without decreasing activation length on the absolute difference between the standardized DNA-1 and standardized non-binding class activation sums [Eq.1]. Here, x and y are activations of DNA-1 and non-bind, respectively. Peak activations were then determined using operator set thresholds.

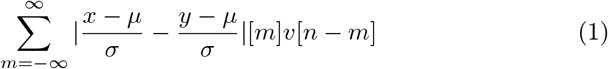

#### 3.6.1 Knockout Test

To validate the activations provided by the hidden state’s analysis on DNA-1, a “knockout” test was performed. We reasoned that if the suspected binding site is being used for class prediction by the model, once such information is removed, the model should be more likely to predict non-binding. For each nonbinding sequence, the residues at the literature binding sites were transplanted into a copy of the DNA-1 sequence at the literature binding site positions. Sequences in both classes were paired according to similar lengths. For binding sequences that were paired with a non-binding sequence of shorter length, a random non-binding sequence was chosen to fill in the binding sites exceeding the original sequence length. Only five non-binding sequences were smaller than the last bind site position, 327. Modified DNA-1 sequences were reversed and evaluated by the top trained prediction models. Hidden states were extracted and processed according to equation one between knockout and nonbinding class activations and compared to DNA-1. To reduce “noise” between major peaks found in both DNA-1 and knockout activation outside the literature binding sites, the difference between DNA-1 and knockout greater than 0 provided an alternative bind site prediction for peaks at various operator-set thresholds.

#### 3.6.2 Insertion Test

To determine if binding sites are all that is necessary for class prediction, this process was repeated for an “insertion” test. DNA-1 literature bind sites were transplanted into non-binding sequences of similar length. If a non-binding sequence was shorter than 444 residues, the length of DNA-1, only the available binding sites were swapped and lengths were retained. Sequences were reversed and evaluated by the top trained models. Hidden states were extracted and processed according to equation one between insertion sequences’ activation and non-bind sequences’ activation. The insertion activation was then compared to the DNA-1 activation and literature known binding sites.

#### 3.6.3 Peak Knockout Test

The knockout test described previously relies on literature binding site knowledge. The work proposed here is attempting to provide viable suggestions for proteomic interactions in cases where domain knowledge is extremely limited, as is often the case for synthetic protein design. As a method of predicting binding sites in such cases, another knockout test was performed on the major peaks in DNA-1 activations (i.e. “peak knockout”). The literature known binding site for DNA-1 is 66 residues, approximately 15% of the total sequence. Therefore, to make balanced comparisons with the original knockout test, peaks above 58% threshold resulted in 68 residues to be modified in the subsequent test. Similar thresholds were chosen for the smaller LSTM and CNN model variants. Positions of these peaks were then used as the sites that were replaced by non-binding sequence residues. All sequences were evaluated accordingly with the original knockout test procedure. Comparisons were then made between the peak knockout and DNA-1 activations. To reduce noise in the activations, the difference between DNA-1 and peak knockout activations yield an alternative binding site suggestion. Final binding site sub-sequences were found by overlapping the activations created by previous DNA-1 analysis peaks and peak knockout occluded DNA-1 activations peaks.

## 4 Results

We found no statistically significant differences in amino acid frequency or first amino acid occurrence. Lengths in the dataset were significantly different resulting in the preprocessing techniques described above [Fig. 4]. TSNE on raw sequences was inconclusive [Fig. 5]. No clusters within classes could lead to insightful inference in function from primary structure provided by the dataset.

**Fig. 4.**
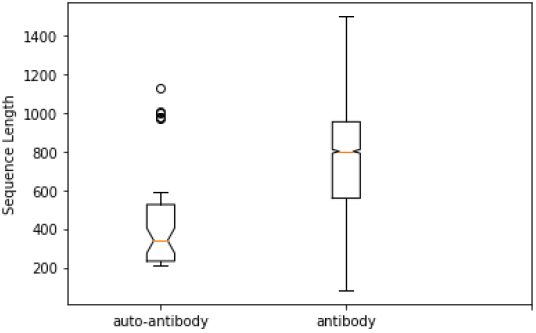
Sequence Length Per Class.

**Fig. 5.**
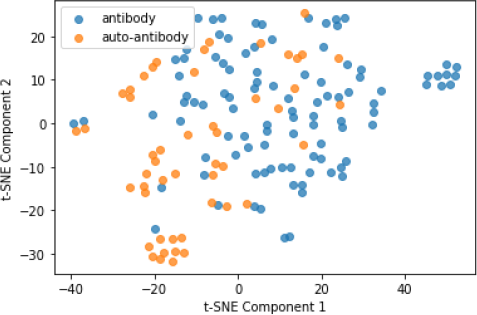
TSNE on raw sequence data.

### 4.1 Binding Specificity

Table One shows the accuracies for each of the four models [Fig. 6]. Across both LSTMs and CNNs, the smaller variants resulted in better accuracy. The smaller CNN model had the best average validation accuracy across 100 models was very statistically significant at 87.81% by one sample t-test, t(99) = 90.9673, p<.0001. The smaller LSTM binding prediction achieved average validation accuracy of 72.64% and was statistically significant, t(99) = 20.5201, p<.0001. Notably, the LSTM prediction performance increases with added sigmoid activation, while the same effect is not observed for CNN. During the testing phase on the trained models, average accuracy score was 87.07% for the binding class and 88.56% for the non-binding class by the LSTM and 96.56%, 97.81% by CNN, respectively for each smaller variant and similarly observed for larger variants. The chosen top models for hidden state analysis were all above 95% accurate on all sequences. PCA on the hidden states were inconclusive [Fig. 7]. Raw activation weight visualization showed distinct horizontal bands at particular time-steps [Fig. 8]. Activation sums showed distinct peaks at different positions per class.

**Fig. 6.**
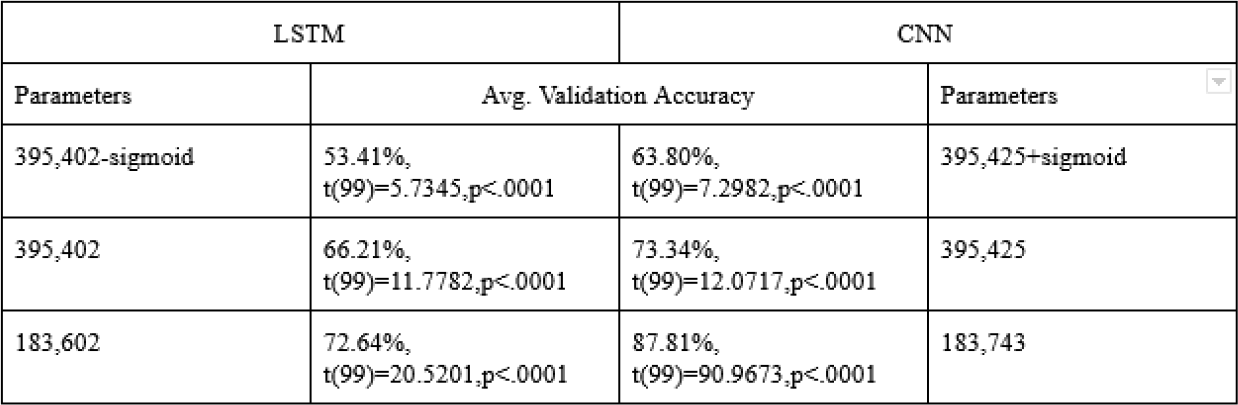
Average validation accuracies across LSTM and CNN models.

**Fig. 7.**
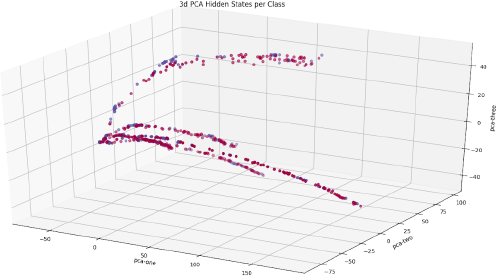
PCA on Hidden Cell Weights. Each hidden cell activation matrix is encoded per sequence sample for binding (red) and non-binding (purple) classes.

**Fig. 8.**
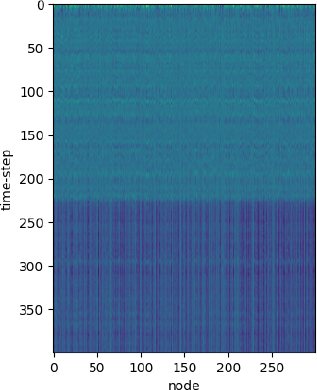
Raw Activation Differences Between Classes. Difference in hidden cell weights from LSTM (left) and feature maps from CNN (right) for all combined bind and non-bind sequences.

### 4.2 Binding-Site Analysis

Table Two shows correlation and significance between the suggested binding sites at each stage of processing for respective models [Fig. 9]. The larger LSTM achieved the best binding site recovery and is reported in the proceeding results. Similar results and figures were observed in other model variants. DNA-1, binding, and non-binding activations had different unique peaks across different positions [Fig. 10]. DNA-1 literature-defined binding site and hidden state activation sums overlapped in several major peaks [Fig. 11] but were not significantly correlated, r(798) .0627, p > .1. Processed DNA-1 activations according to equation one, r(798) = .1870, P < .001 and subsequent peaks at threshold 85%, r(798) = .2476, p < .0001, and threshold 58%, r(798) = .1433, p < .01, were significantly correlated [Fig. 12]. Knockout testing results showed on average 86.17% reversal from binding to non-binding class prediction. Hidden state analysis showed similar general trends between the knockout and the DNA-1 activations and was not significant, r(798) = −.0243, p > .1. At threshold 58%, however, trending significance was observed between knockout and literature known sites, r(798) = −.0903, p < .1. Discrepancies in activation difference between knockout and DNA-1 were least in areas outside the literature binding sites. Thus, occlusion of knockout activations from DNA-1 activations shows significant noise reduction between literature known binding sites, r(798) = .2140, p < .0001. Subsequent peaks for threshold 58% was not significant, r(798) = .0328, p > .1. However, lowering this threshold to 50% resulted again in significance between the knockout occluded DNA-1 activations and literature binding sites, r(798) = .1122, p = .02. Insertion test was less effective showing only 6.67% reversal on average with upper bounds at 9.87%. Hidden state analysis showed insertion activations very closely following DNA-1 activations, but without significant correlation to the binding sites, r(798) = .0500, r > .1. However, only looking at peaks above 58% threshold did show significance, r(798) = .1018, p =.04. Knockout of all DNA-1 peaks greater than 58% the max peak [Fig. 12] showed on average 80.74% prediction reversal, with three models correctly predicting 80 or more sequences. Hidden states analysis was similar to that of previous knockout, with activations closely following DNA-1 activations with most discrepancies inside literature binding regions resulting in non-significance, r(798) = −.0415, p > .1. Peaks above 58% threshold were trending towards significant, r(798) = −.0880, p < .1 [Fig. 13]. Noise reduction by difference between DNA-1 and knockout peak activations showed significant overlap between the literature known binding sites and the model suggested sites peaks at 58% threshold, r(798) = .2566, p < .0001 [Fig. 14].

**Fig. 9.**
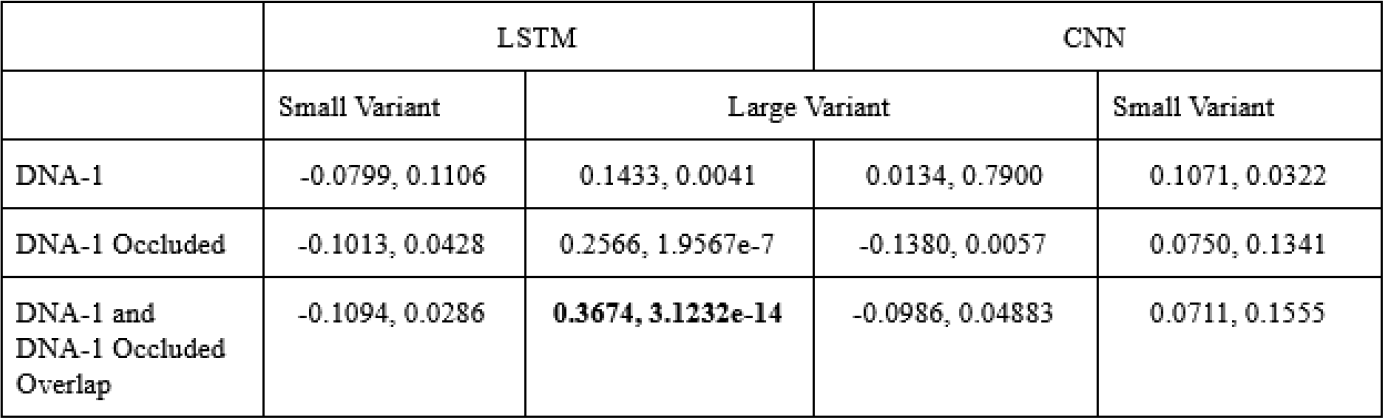
Pearson correlation coefficients and significance values for binding site activation’s at various processing steps compared to literature bind site for DNA-1, df=798.

**Fig. 10.**
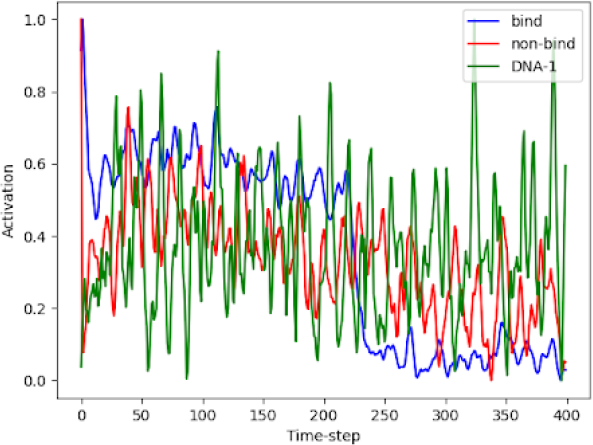
Non-binding, binding, and DNA-1 activations from LSTM hidden cell standardized between 0 and 1.

**Fig. 11.**
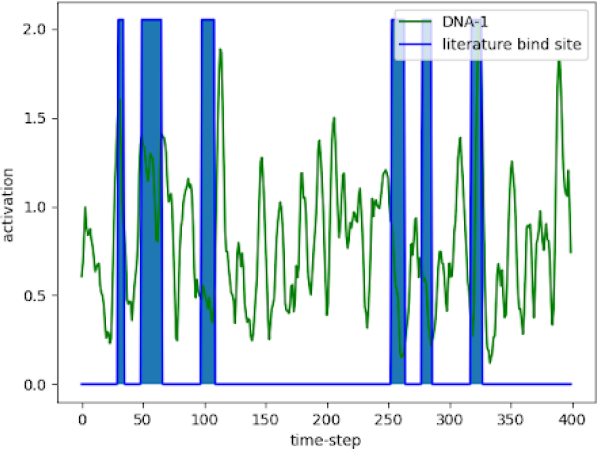
DNA-1 standardized activations before equation one processing, r(798) = 0.053.

**Fig. 12.**
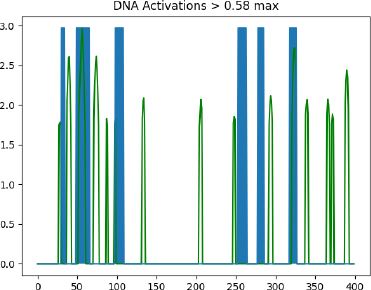
DNA-1 activations processed according to equation one for peaks at 58% threshold, r(798) = 0.143).

**Fig. 13.**
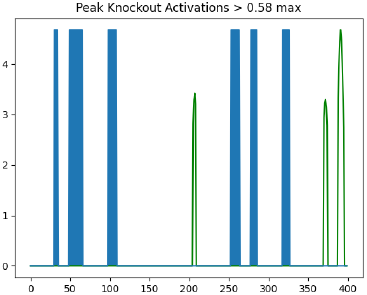
Knockout of DNA-1 peaks processed according to equation one for peaks at 58% threshold, r(798) = −0.088.

**Fig. 14.**
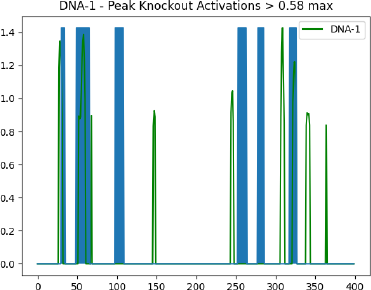
Peak knockout occluded DNA-1 activations processed according to equation on for peaks at 58% threshold, r(798) = 0.257.

### 4.3 Binding sub-sequences

Recovered sub-sequences found only by the 58% threshold peaks of DNA-1, peak knockout procedure (without overlap) recovered significant partial residues in half the literature binding regions, r(798) = .256, p < .0001 [Fig. 15]. In figures 15 and 16, red letters indicate model suggested regions of interest, bold as literature suggested binding residues, and underlined regions indicating correctly suggested residues. Six other residues and sub-sequences outside the literature binding sites were also suggested. There is no apparent trend in the nature of residues suggested in blatant misses. However, some near-hits are often only a few residues premature of literature bind site subsequences. Overlap between DNA-1 peaks and peak knockout occluded DNA-1 peak activations resulted in highest subsequence fidelity in significance, r(798) = .367, p < .0001 [Fig. 16]. This final method resulted in shorter and less frequent misses. Only one region that was originally a hit was missed, however, a near miss suggestion was directly next to the target residues in this site. Agreement between CNN and LSTM binding sub-sequences is shown in figure 17.

**Fig. 15.**
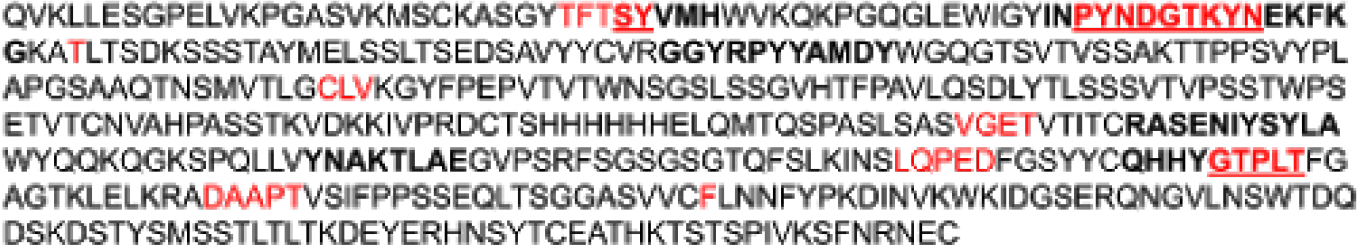
Sub-sequences found in knockout peak analysis. Peaks according to non-bind occluded DNA-1 activation peaks at 58% threshold and model suggested peaks via knockout peak occluded DNA-1 activations where red indicates model suggested sites, bold is literature binding sites, and underline is the overlap of the two.

**Fig. 16.**
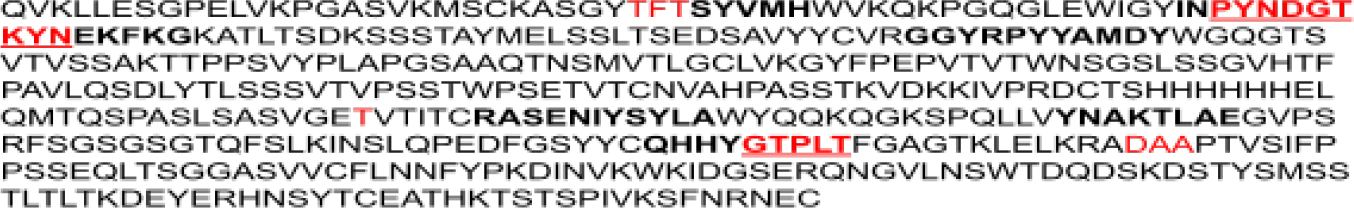
Overlap of DNA-1 and difference between DNA-1 with peak knockout peaks at 58% and sub-sequence comparison with literature binding site where red indicates model suggested sites, bold is literature binding sites, and underline is the overlap of the two.

**Fig. 17.**
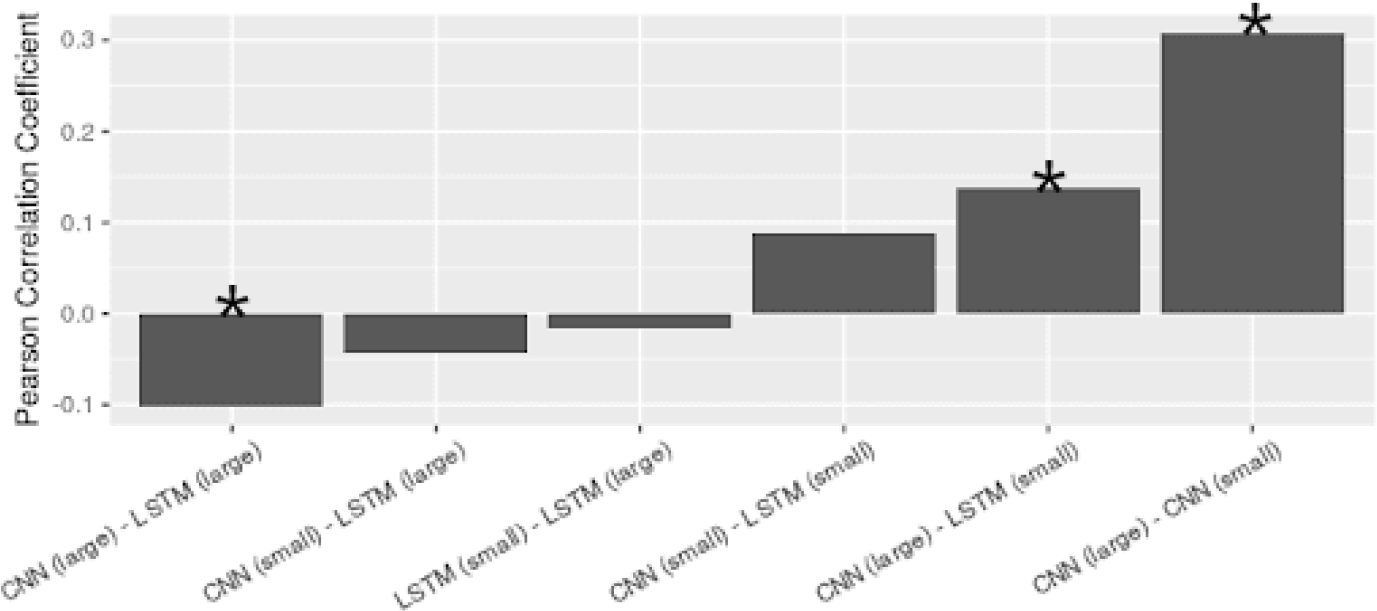
Agreement between model variants on subsequence prediction. From left to right: r(798) = −0.1018, p < .05; r(798) = −.0442, p > .1; r(798) = −.0155, p > .1; r(798) = .0892, p < .1; r(798) = .1369, p < .01; r(798) = .3070, p<.001.

## 5 Discussion

Lack of structural/functional properties separable by TSNE embedding space and brute-forced attributes of raw data shows the need for complex feature representations offered by deep learning models.

### 5.1 Binding Classification

As expected, model architecture configurations with less trainable parameters performed better. Since the optimizable gradients are less complex, the smaller dataset used in this application is more thoroughly integrated during back propagation. Overall, CNN models perform better on prediction tasks. Hyperparameters for both types of models drastically change performance. Thus, a combination of both types of models could best aid in a more vigorous approach to vaccine and drug design.

Model variability in training is due to differences in random weight initialization and order of batch sampling creating hurdles in overcoming localminima during parameter optimization. Binding prediction has slight bias for non-binding class, supporting the absence of overfitting to the binding dataset. Models with higher test accuracy, even if they didn’t perform as well during the training phase, show accurately learned weights for class recognition, which can be extended to novel or synthetic proteins outside the training and validation dataset. Activation differences across all nodes at specific positions suggest model decisions are portrayed differently per class in the hidden weights. Furthermore, the raw activation across nodes shows the model relies most on sequence positions up to 225 while following positions are less important for binding class prediction. Since most sequences are around this length or greater than, this supports the HVD region, at the beginning of the FASTA, being the most common region of binding. These areas of interest are further shown in the activation sums which lead to the sub-sequence distribution.

### 5.2 Binding Site Analysis

Hidden state analysis showed greater correspondence to previously established binding sites in the LSTM vs. CNN models. This is likely because LSTM encodes position specific information rather than CNN which detects spatial-temporal invariant features. CNN top models suffered in their ability to correctly predict the sequence of interest, DNA-1. This likely resulted in poorly interpreted hidden state analysis, supported by figure 17. Furthermore, the overall increased PC for larger models suggests that while binding prediction is increased with less parameters, the learned features for extracting binding sites are more interpretable in models with more parameters. Difference between DNA-1 and the overall bind activation weights suggests the binding motif is not only position specific but also sub-sequence dependent, otherwise drop-off for later position indices would have been observed in DNA-1 outside the HVD region. All sites had distinguishable overlapping activation peaks. However, there were mafor extraneous activation peaks at non-binding sites primarily in late downstream regions, which can be explained by the LSTM’s implicit higher activation for beginning sequences due to lack of recurrent information. The low PC value in raw activations suggests activation alone is not sufficient enough for high-fidelity binding site suggestions. This is supported by increased PCs from equation one and noise occlusion processing.

#### 5.2.1 Knockout

Majority reversal by knockout test suggests the prediction model relies heavily on literature binding site positions as features for class prediction. Furthermore the PC dropped dramatically for the overall trend and thresholded peaks, suggesting removal of the binding sites impares the models ability to find correct binding sub-sequences, as expected. Discrepancies in peaks between DNA-1and knockout activations were mostly in binding regions. Peaks outside of the literature binding sites, “noise”, were reduced in the occlusion processing step. This intermediate noise is likely caused by high variation amongst training sequences. That is, the model is looking in those positions for learned features it has expected from other sequences during training (ie. feature of proteins in general or type of antibody, etc.). Differences between DNA-1 and knockout suggest these areas are less likely to be true binding site predictions and their removal generally increases PC in lower operator-set thresholds.

#### 5.2.2 Insertion

These assertions are further supported by the weak reversal shown by the insertion of binding sites into non-binding sequences. As insertion test activation trends were most similar to DNA-1 trends and low reversal rates were observed, the model is relying, in part, on other areas of interest due to its training on sequences of various lengths and antibody families. Peaks between the first and last groups of literature binding sites are located where all nodes had activation drop off in the raw visualization. These regions are likely the end of the HVD, proposing the model is also looking for features in this HVD region (residue 0-225) for binding prediction as previously learned in the training phase. While this effect does confound the binding site prediction, we propose it strengthens the overall prediction mechanism’s ability to generalize.

#### 5.2.3 Peak Knockout

Knockout of DNA-1 peaks further support this conjecture as the reversal rate was retained and occlusion of these activations from DNA-1 resulted in the highest significant correlations with literature binding sites. Remarkably, the recovery of binding site information corresponding with literature known binding sites from the peak knockout poses this methodology as reliable for binding site suggestions without available domain knowledge. Making it a unique and helpful technique in synthetic design.

#### 5.2.4 Binding Sites

Subsequence recovery, while somewhat significant without overlap between original DNA-1 peak and peak knockout occluded DNA-1 activations, suggested sites unrelated to binding which could delay research and development. Therefore, the final overlap process shows a highly significant method of computationally predicted residue binding sites and sub-sequences without domain knowledge, limited data infrastructure, and low computing resource requirements. Operators can leverage precision and recall in the binding site suggestion methodology by altering the threshold for peak identification and smoothing operator during convolutions throughout the procedure according to specific use-case needs.

## 6 Conclusion

The current work establishes that the deep learning models applied to primary sequences can predict whether a novel sequence will bind to DNA and that the hidden activations of these models yielded significant agreement with the literature with regard to binding site, r(798) = 0.3674, p = 3.1232e-14. These recovered areas allow researchers to closely examine the network’s internal state; gaining insight into position-specific residue involved in antibody:DNA-binding. We also show while CNN is better suited for binding prediction in smaller models, larger LSTM hidden states allow for a more accurate binding site interpretation. The proposed methodology can be extended to other domains of interest that may have limited datasets available. Future work should focus on reducing noise in the hidden state activations and compiling residue investigations/predictions in a comprehensive manner to inform binding site prediction with end-use researchers in mind. Findings implicate suggestions for RVD and possible synthetic components. Collective implications of this research will further the rapidly developing field of applied deep learning, which in turn will allow for more efficient applied applications and directly enhance gene and protein data processing. Additionally, we expect the proposed model to be versatile at evaluating other proteomic datasets and user friendly for researchers without extensive computational background knowledge and computing resources. At the same time, the prospective sequence specificities allow experts in wet-lab approaches, like x-ray crystallography, to make more informed decisions.

## Acknowledgements

The authors thank PhD candidate Paul Morris at the Center for Complex Systems and Brain Sciences for their insightful discussions on natural language processing and data analysis.

## Declarations

### 6.3 Availability of data and material

Associated data and material can be found at https://github.com/mpcrlab/AntibodyBindingPrediction.

### 6.4 Code availability

Associated software and code can be found at https://github.com/mpcrlab/AntibodyBindingPrediction.

## References

1. S. Aotsuka, The Ryumachi 28, 96 (1988)

2. M. Teodorescu, Clinical and Applied Immunology Reviews 2(2), 115 (2002)

3. J.A. Beckingham, J. Cleary, M. Bobeck, G.D. Glick, Biochemistry 42(14), 4118 (2003)

4. L. Rodkey, G. Gololobov, C. Rumbley, J. Rumbley, D. Schourov, O. Makarevich, A. Gabibov, E. Voss, Applied biochemistry and biotechnology 83(1-3), 95 (2000)

5. P.C. Swanson, C. Ackroyd, G.D. Glick, Biochemistry 35(5), 1624 (1996)

6. L. Spatz, A. Iliev, V. Saenko, L. Jones, M. Irigoyen, A. Manheimer-Lory, B. Gaynor, C. Putterman, M. Bynoe, C. Kowal, et al., Methods 11(1), 70 (1997)

7. M. Pavlovic, R. Chen, A.M. Kats, M.F. Cavallo, S. Saccocio, P. Keating, J.X. Hartmann, Annals of the New York Academy of Sciences 1108(1), 203 (2007)

8. A. Kozyr, IUBMB Life 39(2), 403 (1996)

9. J.N. Herron, X. He, D. Ballard, P. Blier, P. Pace, A. Bothwell, E. Voss Jr, A. Edmundson, Proteins: Structure, Function, and Bioinformatics 11(3), 159 (1991)

10. J.J. Tanner, A.A. Komissarov, S.L. Deutscher, Journal of molecular biology 314(4), 807 (2001)

11. D. Gu, Y. Zhou, V. Kallhoff, B. Baban, J.J. Tanner, D.F. Becker, Journal of Biological Chemistry 279(30), 31171 (2004)

12. Z. Ou, C.A. Bottoms, M.T. Henzl, J.J. Tanner, Journal of molecular biology 374(4), 1029 (2007)

13. M. Pavlovic, A. Kats, M. Cavallo, Y. Shoenfeld, Lupus 19(7) (2010)

14. M. Pavlovic, (2009)

15. O. Russakovsky, J. Deng, H. Su, J. Krause, S. Satheesh, S. Ma, Z. Huang, A. Karpathy, A. Khosla, M. Bernstein, et al., International journal of computer vision 115(3), 211 (2015)

16. A. Radford, K. Narasimhan, T. Salimans, I. Sutskever. Improving language understanding by generative pre-training (2018)

17. A.W. Senior, R. Evans, J. Jumper, J. Kirkpatrick, L. Sifre, T. Green, C. Qin, A. Žídek, A.W. Nelson, A. Bridgland, et al., Nature 577(7792), 706 (2020)

18. A. Trabelsi, M. Chaabane, A. Ben-Hur, Bioinformatics 35(14), i269 (2019)

19. S. Min, B. Lee, S. Yoon, Briefings in bioinformatics 18(5), 851 (2017)

20. B. Alipanahi, A. Delong, M.T. Weirauch, B.J. Frey, Nature biotechnology 33(8), 831 (2015)

21. X. Pan, H.B. Shen, BMC bioinformatics 18(1), 136 (2017)

22. X. Pan, P. Rijnbeek, J. Yan, H.B. Shen, BMC genomics 19(1), 511 (2018)

23. M. Paul, S.C. Rachel, E.H. William, B. Elan, Journal of chemical information and modeling

24. A. Rives, S. Goyal, J. Meier, D. Guo, M. Ott, C.L. Zitnick, J. Ma, R. Fergus, bioRxiv p. 622803 (2019)

25. S. Wang, Y. Guo, Y. Wang, H. Sun, J. Huang, in Proceedings of the 10th ACM International Conference on Bioinformatics, Computational Biology and Health Informatics (2019), pp. 429–436

26. T. Sun, B. Zhou, L. Lai, J. Pei, BMC bioinformatics 18(1), 1 (2017)

27. X. Liu, arXiv preprint arXiv:1701.08318 (2017)

28. Z. Zhao, X. Gong, IEEE/ACM transactions on computational biology and bioinformatics (2017)

29.x J. Liu, X. Gong, BMC bioinformatics 20(1), 609 (2019)

30. S. Hochreiter, J. Schmidhuber, in Advances in neural information processing systems (1997), pp. 473–479

31. U. Consortium, Nucleic acids research 47(D1), D506 (2019)

32. S. Pietrokovski, S. Henikoff, Molecular and General Genetics MGG 254(6), 689 (1997)

33. T. Hou, K. Chen, W.A. McLaughlin, B. Lu, W. Wang, PLoS Comput Biol 2(1), e1 (2006)

34. M. Nielsen, C. Lundegaard, O. Lund, BMC bioinformatics 8(1), 238 (2007)

35. C. Mooney, G. Pollastri, D.C. Shields, N.J. Haslam, Journal of molecular biology 415(1), 193 (2012)

36. S.H. Yoon, S.M. Ha, S. Kwon, J. Lim, Y. Kim, H. Seo, J. Chun, International journal of systematic and evolutionary microbiology 67(5), 1613 (2017)

37. H. Berman, J. Westbrook, Z. Feng, G. Gilliland, T. Bhat, H. Weissig, Shindyalov, and PE Bourne pp. 235–242 (2000)

